# Mechanism of the Covalent Inhibition of Human Transmembrane Protease Serine 2 as an Original Antiviral Strategy

**DOI:** 10.1101/2023.04.23.537985

**Authors:** Angelo Spinello, Luisa D’Anna, Emmanuelle Bignon, Tom Miclot, Stéphanie Grandemange, Alessio Terenzi, Giampaolo Barone, Florent Barbault, Antonio Monari

**Author notes:** AUTHOR INFORMATION **Corresponding Author** F.B., A.M.

## Abstract

The Transmembrane Protease Serine 2 (TMPRSS2) is a human enzyme which is involved in the maturation and post-translation of different proteins. In addition of being overexpressed in cancer cells, TMPRSS2 plays a further fundamental role in favoring viral infections by allowing the fusion of the virus envelope and the cellular membrane, notably in SARS-CoV-2. In this contribution we resort to multiscale molecular modeling to unravel the structural and dynamical features of TMPRSS2 and its interaction with a model lipid bilayer. Furthermore, we shed light into the mechanism of action of a potential inhibitor (Nafamostat), determining the free-energy profile associated with the inhibition reaction, and showing the facile poisoning of the enzyme. Our study, while providing the first atomistically resolved mechanism of TMPRSS2 inhibition, is also fundamental in furnishing a solid framework for further rational design targeting transmembrane proteases in a host-directed antiviral strategy.

**TOC GRAPHICS:** 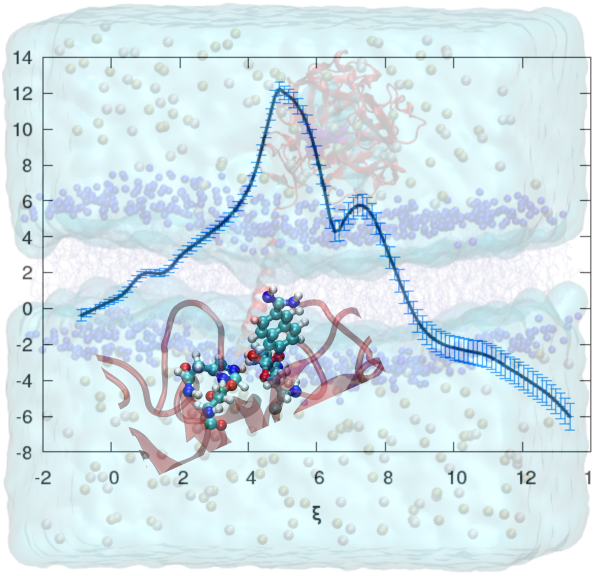

---

The transmembrane protease serine 2 (TMPRSS2) enzyme belongs to a family of human proteins^1^ which are anchored to the cellular membrane and participate to various protective and regulative pathways. From a biochemical point of view, TMPRSS2 exerts a proteolytic activity, which is needed to assure the degradative remodeling of the extracellular medium^2^ and the proteolytic activation of different membrane proteins.^3^ Interestingly, TMPRSS2 and the related proteins are also involved in key epithelial homeostatic regulation.^4^ Furthermore, TMPRSS2 is highly overexpressed in different cancer types, notably prostate, invasive breast, head and neck cancers, and lung carcinoma, making it an attractive biomarker for the early diagnosis.^5–7^ TMPRSS2 is related to carcinogenesis, cancer metastatic progression,^8,9^ and its role on the chronic pain experienced by patients has also been stressed out.^10^ Indeed, its localization at the surface of cancer cells allows the protein to mediate, via its proteolytic activity, different signaling cascades and transduction between the cells and the extracellular medium, including pain signals. However, the full mechanisms relating TMPRSS2 to cancer induction and progression, as well as its role in pain sensitization, are neither known at cellular level nor at chemical one. The involvement of TMPRSS2 in cancer progression and its link with tumor invasiveness have spurred a significant research activity aimed at identifying potential inhibitors.^10^

In addition to its important role in oncology, TMPRSS2 is also fundamental in mediating the response to viral infections.^11–17^ Indeed, while not acting as the main cellular receptor, TMPRSS2 is necessary to allow the fusion of the viral and cellular membranes, hence to the entry of the viral genetic material which triggers infection. As a matter of fact, this role of TMPRSS2 has been particularly stressed in the case of SARS-CoV-2, i.e. the causative agent of the COVID-19 pandemics.^18^ Due to the high societal, sanitary, and economics burdens brought by SARS-CoV-2, and more generally emerging infectious diseases, particularly RNA-viruses, the importance of TMPRSS2 can hardly be overestimated, as well as the beneficial development of inhibitors which would constitute valuable assets to deploy a rapid pharmacological response, also complementing the vaccinal strategy.

The molecular processes leading to viral infection,^19,20^ and the related hijacking of the cellular functions to assure the virus fitness and reproduction, are different and complex, and are regulated by a number of structural and non-structural proteins. The latter interfere with different host functions, including autophagy,^21,22^ immune system,^23–25^ or membrane polarization,^26,27^ while permitting the replication of the viral genome, its assembling into nascent virions, and finally the infection of new cells. In the case of SARS-CoV-2, which is an enveloped, positive-strand RNA virus belonging to the family of beta-coronaviruses, the infection of host cells is mediated by the interaction of the Spike (S) protein^28,29^ present on the viral envelope, with a cellular receptor, the angiotensin-converting enzyme 2 (ACE2), which is a transmembrane enzyme involved in the regulation of blood pressure, and densely present on the surface of lung and intestinal cells. The interaction between S, which is a large trimeric protein, and ACE2 proceeds via the formation of extended hydrogen-bonds networks with the receptor-binding domain (RBD),^30–33^ which in turns induce a conformational transition allowing the membrane fusion. However, to complete the fusion, and hence to develop the viral infection, the cleavage of ACE2-bound S at its furin cleavage site is necessary.^34,35^ It has been shown that the absence of furin cleavage sites in S suppresses its infective capacity, notably in ferrets.^36^ The cleavage of S, upon binding to ACE2, is exerted by TMPRSS2, which, thus, should be regarded as a key factor allowing and facilitating SARS-CoV-2 infection.^37–39^ Hence, its inhibition would represent a most suitable therapeutic strategy, which could complement not only vaccine design, but also the virus-targeting drugs including protease and polymerase inhibitors.

From a structural point of view^11^ TMPRSS2 is a rather large enzyme, consisting of an intrinsically disordered intracellular domain, an anchoring transmembrane helix, and an extended structured extracellular domain on which the protease active site is located. The extracellular domain can be further subdivided into three subdomains, having different functions: a low density lipoprotein receptor, one class A scavenger receptor cysteine-rich area, and finally the catalytic subdomain, which is located in the C-terminal region of the protein.^11^ The proteolytic activity is exerted through a catalytic triad comprising His296, Asp345, and Ser441, which is located at the periphery of the protein in a solvent exposed pocket, allowing a relatively easy access to the substrate.^11^ The structural stability of the protein, and in particular the mutual orientation of the extracellular subdomains, is also reinforced by the presence of an extended network of disulfide-bridges, between spatially closed cysteines.^11^

Because of the function of TMPRSS2, and of its low selectivity allowing for the cleavage of different substrates, its covalent inhibition is particularly promising for therapeutical purposes. To this end, among the different inhibitor compound proposed, Nafamostat (NUF) ^11^ forms esters with the catalytic serine upon an irreversible transesterification reaction, thus irreversibly blocking the catalytic activity of the enzyme.

In the present contribution, we have modeled the structure and dynamics of TMPRSS2 embedded in a model lipid bilayer. The structure of TMPRSS2 has been obtained from the recent experimental structure published by Fraser at al. (pdb code: 7meq).^11^ The missing residues were inserted using SwissModel,^40^ while the disordered intracellular domain and the covalently-bound NUF were deleted. TMPRSS2 was embedded in a modeled lipid bilayer consisting of 415 1-palmitoyl-2-oleoyl-sn-glycero-3-phosphocholine (POPC) lipids. The initial system was built using the charmm-gui web interface^41,42^ and a buffer of water was added including KCl ions to reach a physiological concentration of 0.15 M (see Figure 1A). After preliminary minimization and equilibration, by progressively removing positional constraints on the lipids and protein backbone atoms, molecular dynamic (MD) simulations have been performed for about 500 ns in the isothermal and isobaric ensemble (NPT) by using the NAMD code.^43,44^ Once the protein equilibrated, the NUF inhibitor has been re-docked to the protein using AutoDock,^45^ and a further 100 ns MD simulation has been performed. Finally, the non-covalent binding free energy profile has been obtained using umbrella sampling as implemented in Amber.^46^ The covalent inhibition of TMPRSS2 by NUF has been assessed by calculating the associated reaction free energy profile, at hybrid QM/MM level. To this end the adaptive string method,^47^ interfaced with amber, has been applied at semiempirical DFTB level,^48^ using the DFTB3 approach^49,50^ and the 3QB parameters.^51^ Further details on the Computational and Methodological strategy are provided in Electronic Supplementary Information (ESI).

**Figure 1.**
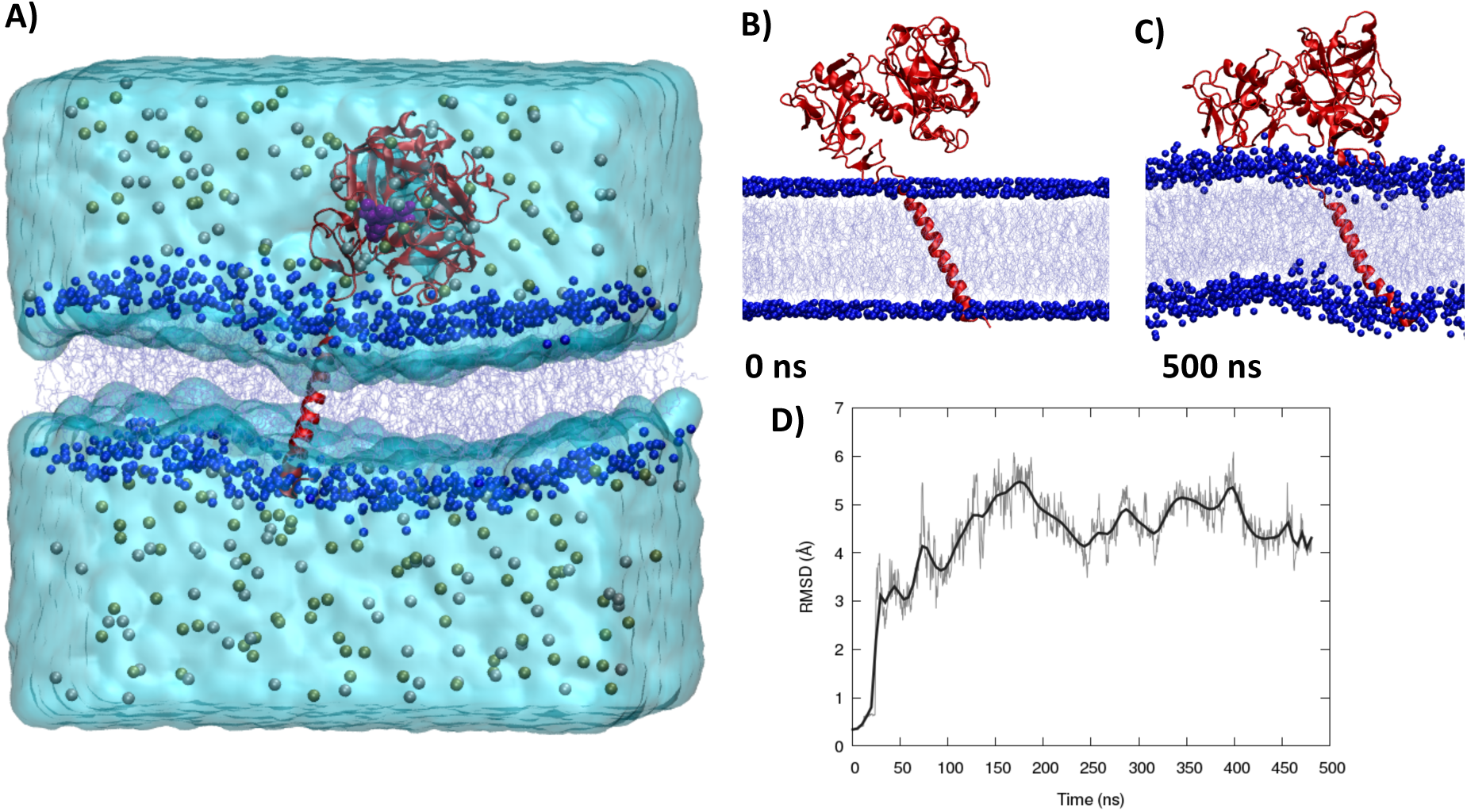
A) The simulated system comprising TMPRSS2 (represented in red cartoon) the POPC bilayer (in blue van der Waals and lines) water box (in transparent surface) and KCl counterions. The position of the catalytic triad is evidenced by a solid purple surface. Initial (B) and final (500 ns C) snapshots of the equilibrium dynamics of TMPRSS2 embedded in a lipid bilayer. D) Time series of the root mean square deviation (RMSD) of TMPRSS2 along the equilibrium MD simulation.

As can be appreciated in Figure 1, TMPRSS2 is indeed forming a stable aggregate with the POPC membrane, optimizing protein-lipid interactions during the simulations. In particular, and as expected, the transmembrane helix persistently inserts into the bilayer, inducing a stable anchoring point. Besides some minor structural rearrangements and some large-amplitude low-frequencies oscillations of the extracellular domain approaching the lipid polar head regions (Figure 1B and 1C), the structural rearrangements of TMPRSS2 are modest. Indeed, the root mean square deviation (RMSD) for the whole protein averages at 4.3 ± 1.0 Å (Figure 1D), further confirming the stability of the complex.

However, the native TMPRSS2 is not catalytic active and, instead, the protein should self-cleave its backbone at the Arg255-Ile256 site to achieve the full enzymatic potentiality.^11,52,53^ For this reason, we have repeated the MD simulation on a system in which the protein backbone was cut at the 255-256 position, thus providing additional C-and N-termini. The results of the cleaved systems are reported in Figure 2. In particular, we can appreciate that, as soon as the positional constraints are removed, the two fragments of the backbone separate, and while the Arg255 terminus is largely solvent-exposed, Ile256 remains closer to the protein surface (Figure 2A-C). As a matter of fact, after the removal of the positional constraints on the protein backbone, the distance between Arg255 carbonyl oxygen and Ile256 terminal nitrogen reaches 40.0 Å to further decreases back to the 30 Å range, indicating a rather uncoupled behavior of the two fragments (Figure 2A).

**Figure 2.**
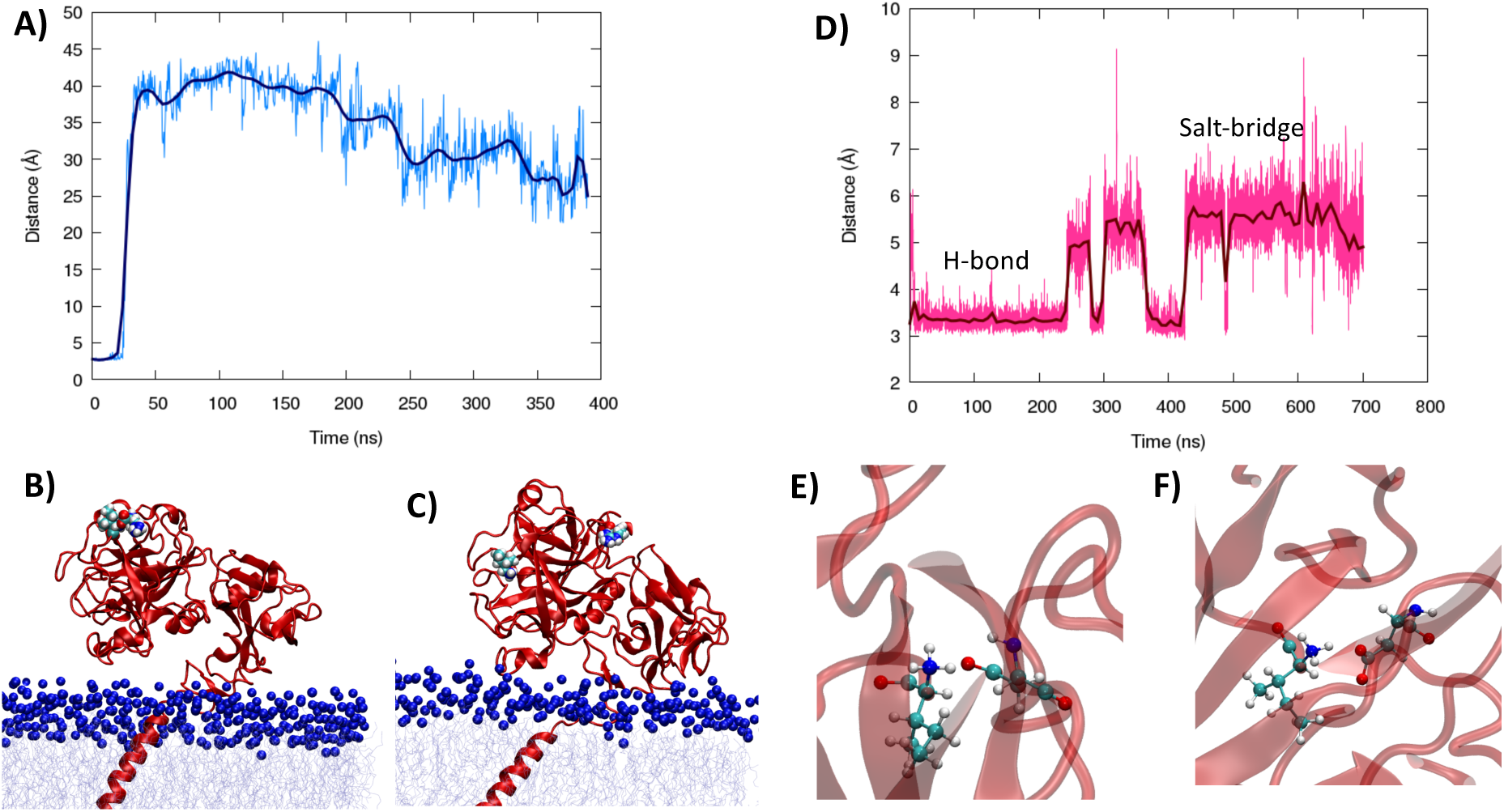
Time evolution of the distance between the oxygen atom of the terminal carbonyl in cleaved residue Arg255 and the terminal nitrogen atom of Ile256 in the equilibrium MD prior to SDM, i.e. the premature conformation (A) and representative snapshots taken at the beginning (B) and at the end (C) of the MD simulation highlighting the separation of Arg255 and Ile256 residues visualized in van der Waals representation. D) Time evolution of the distance between the charged N-terminal nitrogen of Ile256 and the carbon atom in the lateral carboxyl group of Asp440, in the MD simulation following SDM, i.e. the fully mature conformation. The two main conformations driving the Ile256/Asp440 interaction, namely hydrogen bond (E) and salt bridged (F) are also reported.

Upon cleavage, the positively charged terminal nitrogen of Ile256 is reported to interact strongly with the side chain of Asp440, thus participating in shaping and stabilizing the active site conformation.^11^ Our equilibrium MD simulation is not sufficient to retrieve this interaction, due to the presence of an energy barrier partially imposed by the shielding of Asp440 by other protein residues. To overcome this situation and obtain a fully matured conformations we then apply steered MD (SMD) to diminish the Ile256/Asp440 distance by applying an external bias. After releasing the bias-force, the complex between the two amino acids persists over the whole span of the subsequent MD simulation (700 ns, Figure 2D and ESI). However, two distinct conformations may be evidenced, one presenting a distance between the charged N-terminal Ile256 and the Asp440 of less than 4.0 Å, and a second one appearing at later stages of the dynamics in which the two residues are separated by about 6.0 Å (Figure 2D). As evidenced by the representative snapshots reported in Figure 2E, the short distance conformation corresponds to an arrangement in which the carboxylate group of Asp440 is engaged in hydrogen bond with the terminal −NH_3_^+^ of Ile256. Conversely, the longer distance case is still presenting an unspecific salt bridge between the two opposite charged moieties in which the orientational specificity typical of a hydrogen bond is missing and the two units drift slightly further apart. Yet the interaction is globally persistent, confirming the important role of the maturation and the cleavage in fixing the position of Asp440, thus avoiding its occupancy and shielding of the active site.

Once having obtained a fully mature conformation of TMPRSS2, and with the aim to study its interaction with a potential inhibitor, we have redocked NUF to TMPRSS2, and the initial pose obtained has been further relaxed by applying equilibrium MD simulation. Importantly, the docking procedure yielded a pose which is compatible with the one experimentally solved by Fraser et al.,^11^ in which NUF is covalently bound to Ser441, laying inside a pocket in the vicinity of the active site (Figure 3).

**Figure 3.**
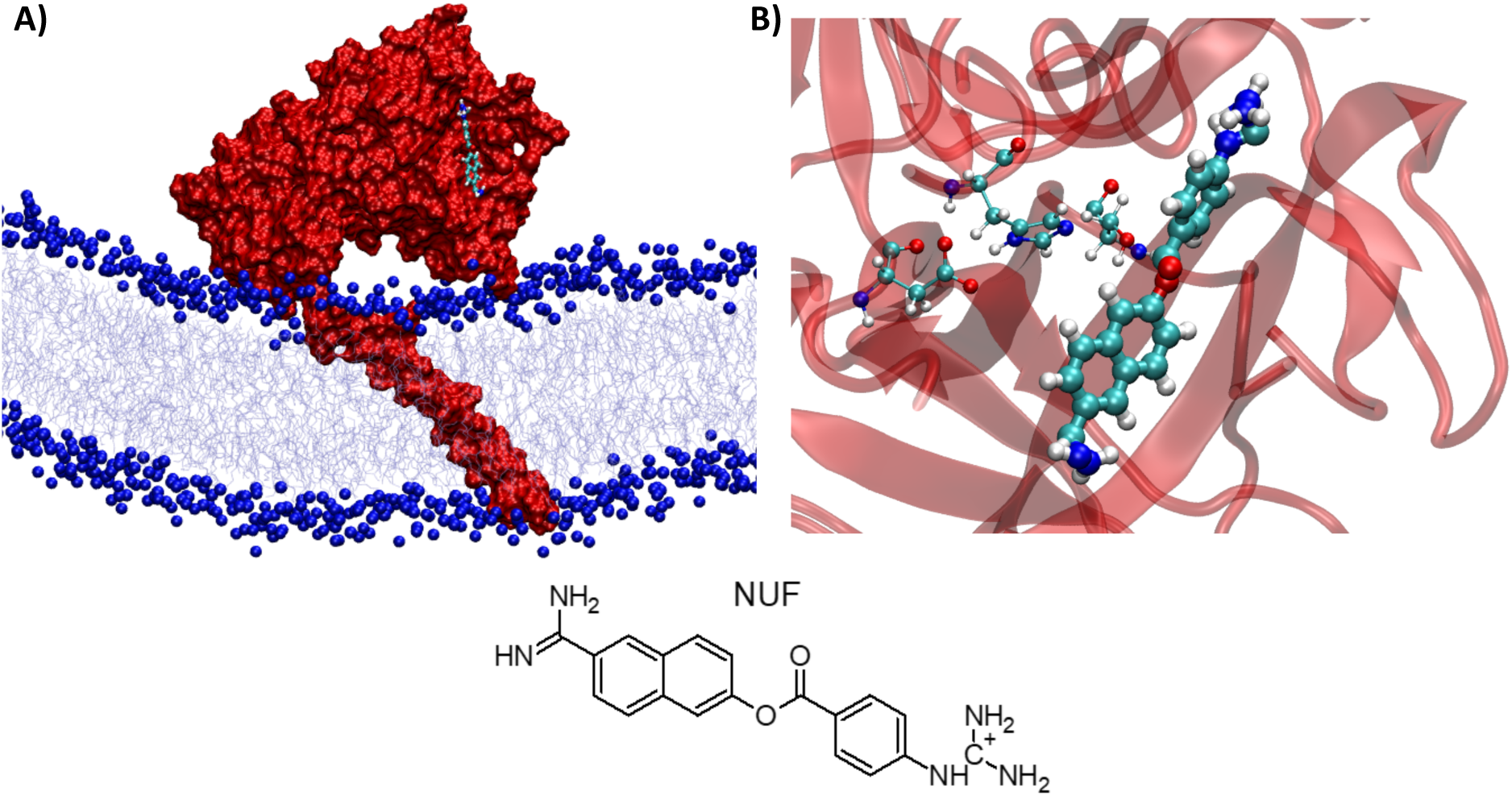
A representative snapshot obtained from the MD simulation after docking of NUF with TMPRSS2 showing the ligand accommodated into the enzyme binding pocket (A). Positioning of NUF with respect to the catalytic triad in the active site (B). The chemical structure of NUF is also given at the bottom.

In particular, the ligand is positioned in proximity to the catalytic triad constituting the active site, and most importantly the OH group of Ser441 is in a favorable orientation to trigger the attack on the carbonyl atom of NUF at a distance of about 8.0 Å. The stabilizing interactions appear mostly driven by non-specific hydrophobic contacts. Yet, some rather labile hydrogen bonds appear along the dynamic such as the one involving the backbone carbonyl of Ala386 and the guanidium groups of NUF.

To better characterize the non-covalent aggregate, we calculated the potential of mean force (PMF) along the dissociation coordinate, by progressively increasing the distance between the center of mass of NUF and of the protein (Figure 4). The dissociation free energy for the non-covalent complex is small and amounts to about 1 kcal/mol. Interestingly, the free energy surface is rather rough and proceeds through different intermediates, in which NUF is occupying alternative positions in the binding pocket. Interestingly, the situation in which NUF is sliding along the surface of TMPRSS2 also corresponds to a minimum free energy. Finally, upon release from the protein, the ligand prefers to interact with the lipid bilayer polar heads, rather than to stay in the water bulk. This behavior is not surprising considering the presence of hydrophobic aromatic rings in the NUF core, flanked with charged peripheral moieties. The PMF reported in Figure 4 clearly confirms that the non-covalent binding of NUF to TMPRSS2 should be extremely labile, and thus not suitable for enzymatic inhibition. Thus, we have also explored the covalent poisoning of TMPRSS2, and more specifically of Ser441. To this aimed, we resorted to QM/MM-based adaptive string method to reduce the dimensionality of conformational space.^47^ While NUF and the side chains of the catalytic triad have been placed in the QM partition, the general collective variable was obtained considering the distance between the lateral oxygen atom of Ser441 and the carbon atom of the carbonyl group of NUF, the ester carbon/oxygen distance in NUF, and finally the distance between the H atom in the hydroxyl group of Ser441 and the deprotonated 𝒳-nitrogen atom of His296 (Figure 5A).

**Figure 4.**
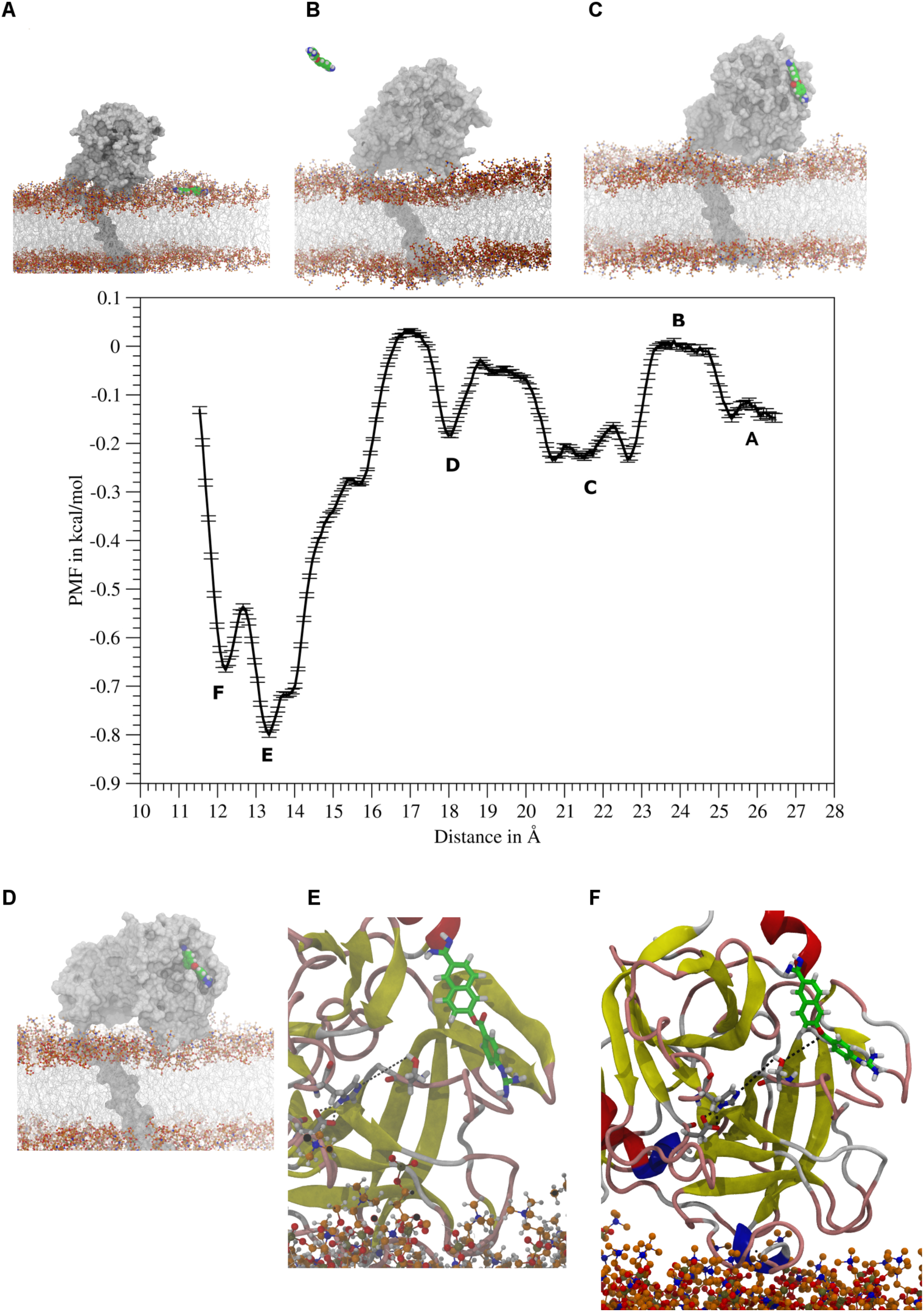
Potential of mean force (PMF) for the separation of the NUF/TMPRSS2 non-covalent complex. Snapshots identifying the most relevant stationary points are also reported.

**Figure 5.**
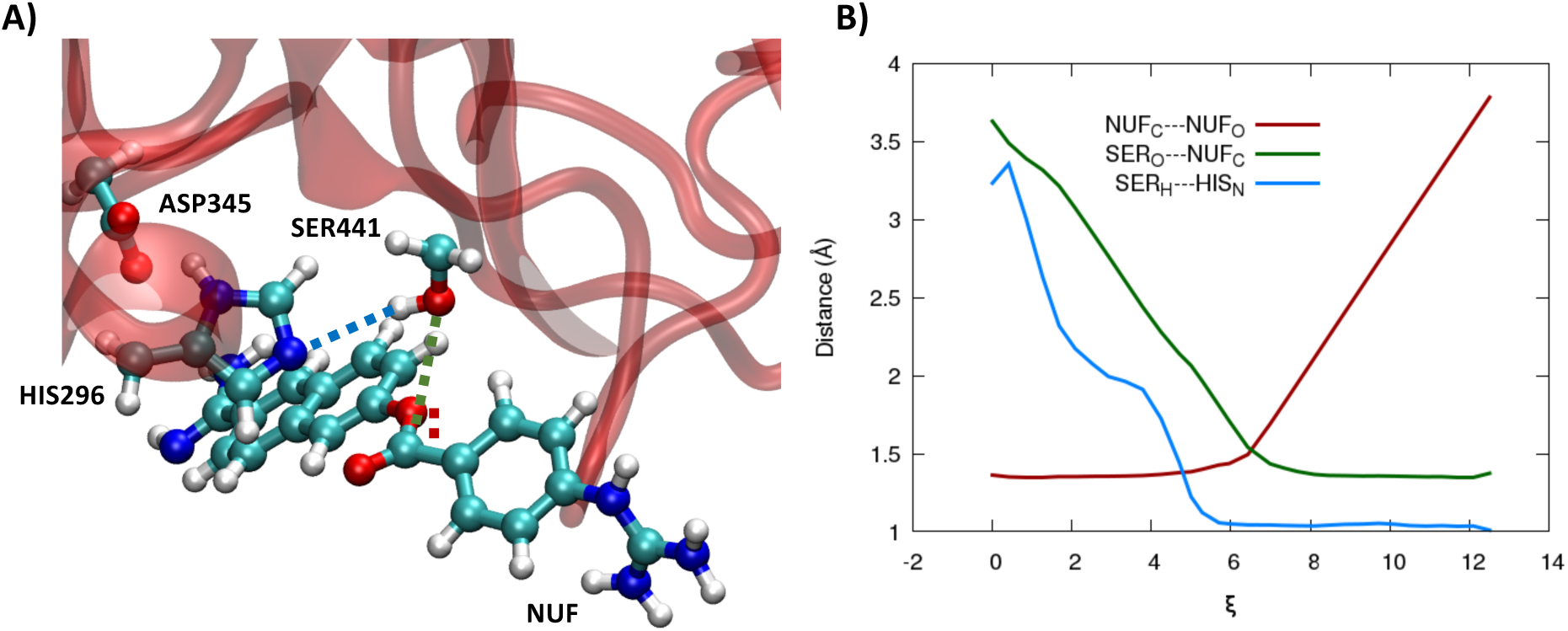
A) QM partition represented in ball and sticks for the QM/MM exploration. The individual distances used to build the string space are indicated in dotted lines. B) Evolution of the individual distances along the minimum energy path in the string space. The same color code as in panel A is used.

The evolution of the individual distances in the optimal string space, corresponding to the minimum energy path is given in Figure 5B. It is evident that the reaction can be characterized as a canonical transesterification, in which first the nucleophilic oxygen of Ser441 attacks the NUF electrophilic carbonyl group, forming a tetrahedral intermediate, while the carbon/oxygen bond originally present in NUF is only subsequently broken, esterifying the catalytic serine. Note that Ser441 is activated by the action of His296 which acts as a proton acceptor for the hydroxyl moiety, making it more nucleophilic. Interestingly, the serine activation slightly precedes the attack to the carbonyl, underlining the sequential character of the mechanism.

The PMF along the optimized string space is reported in Figure 6, along with snapshots showing the structure of critical points on the free energy surface, while the reaction mechanism is schematized in Scheme 1. Interestingly, the overall reaction involves only a rather moderate activation free energy barrier of 12 kcal/mol, while the reaction driving force is of about -6 kcal/mol. Note that this value is slightly underestimated since we have not taken into account the full release of the product, nor the neutralization of the ensuing alcoholate.

**Figure 6.**
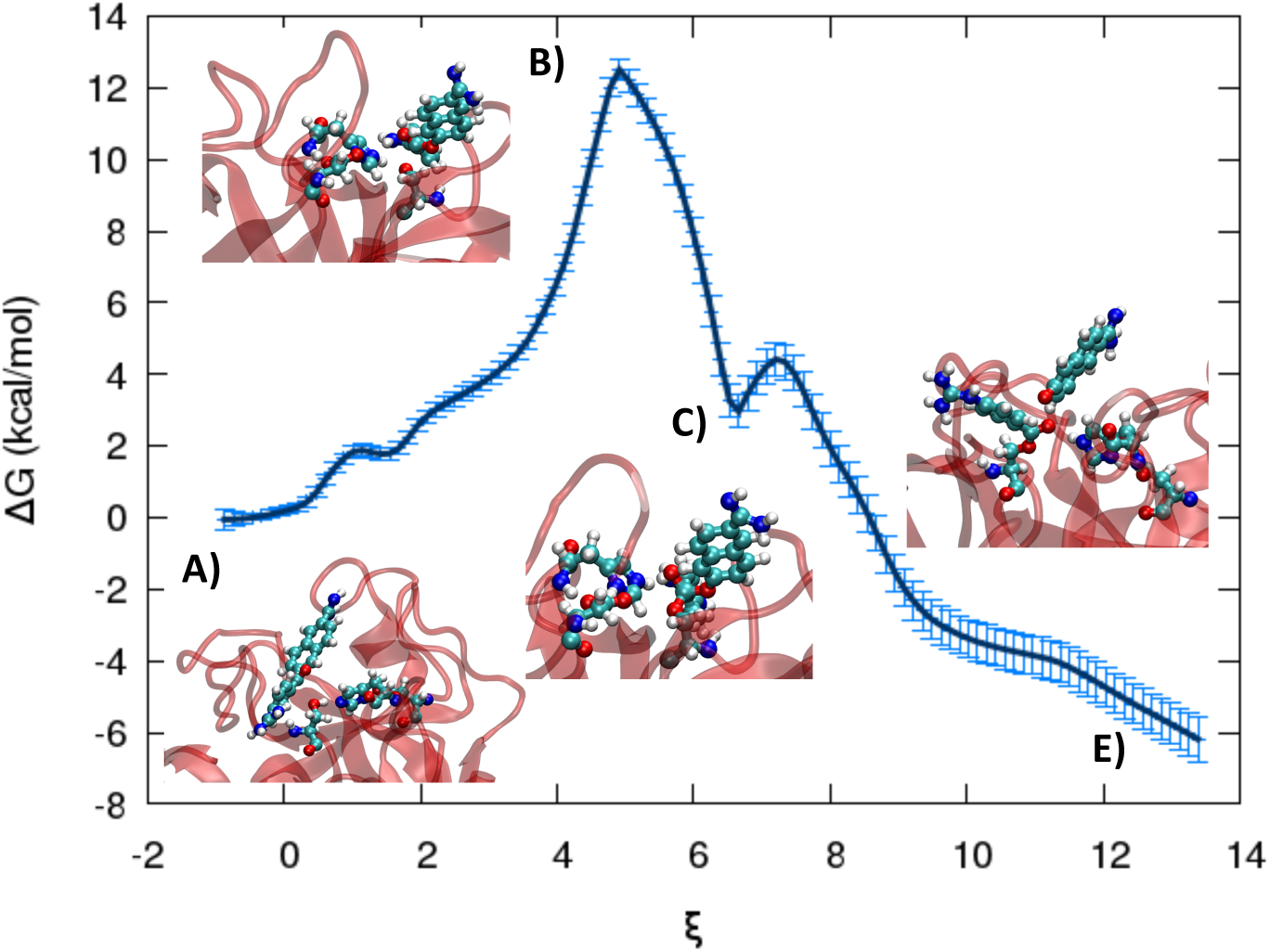
PMF along the optimized string for the reactivity between NUF and TMPRSS2. The snapshots corresponding to the initial (A), transition (B), intermediate (C), and final (E) states are given in inlays and ESI.

Coherently with the transesterification character of the reaction already surmised by the analysis of the optimized string in Figure 5, we may observe the presence of an intermediate (labeled C), which corresponds to the tetravalent sp^3^ hybridized carbon, resulting from the attack of Ser441. A further barrier of less than 2 kcal/mol is necessary to achieve the decomposition of the intermediate, the final esterification of the serine, and the release of the leaving alcoholate. Interestingly, the serine proton appears already fully transferred to the histidine at the transition state, underlining the crucial activation role of this amino acid. This is clearly favored by the presence of the catalytic Asp345, which highly stabilizes the ensuing positively-charged protonated histidine. Finally, the decomposition of the PMF on the individual coordinates shows that the majority of the activation energy is required to form the serine-NUF ester bond (9 kcal/mol, see ESI). On the other hand, the breaking of the NUF carbon-oxygen bond provides the reaction driving force, while the proton transfer has only a very modest barrier of less than 3 kcal/mol, confirming again the crucial role of the catalytic histidine.

Thus, the analyses of the QM/MM simulations lead us to infer the mechanism proposed in Scheme 1. Interestingly, and partially counterintuitively, Ser441 is transferred at the early stages of the reaction to the catalytical His296, and prior to attaining the TS. This feature allows the formation of an alcholate proceeding to the irreversible transesterification via a tetracoordinate carbon intermediate.

**Scheme 1.**
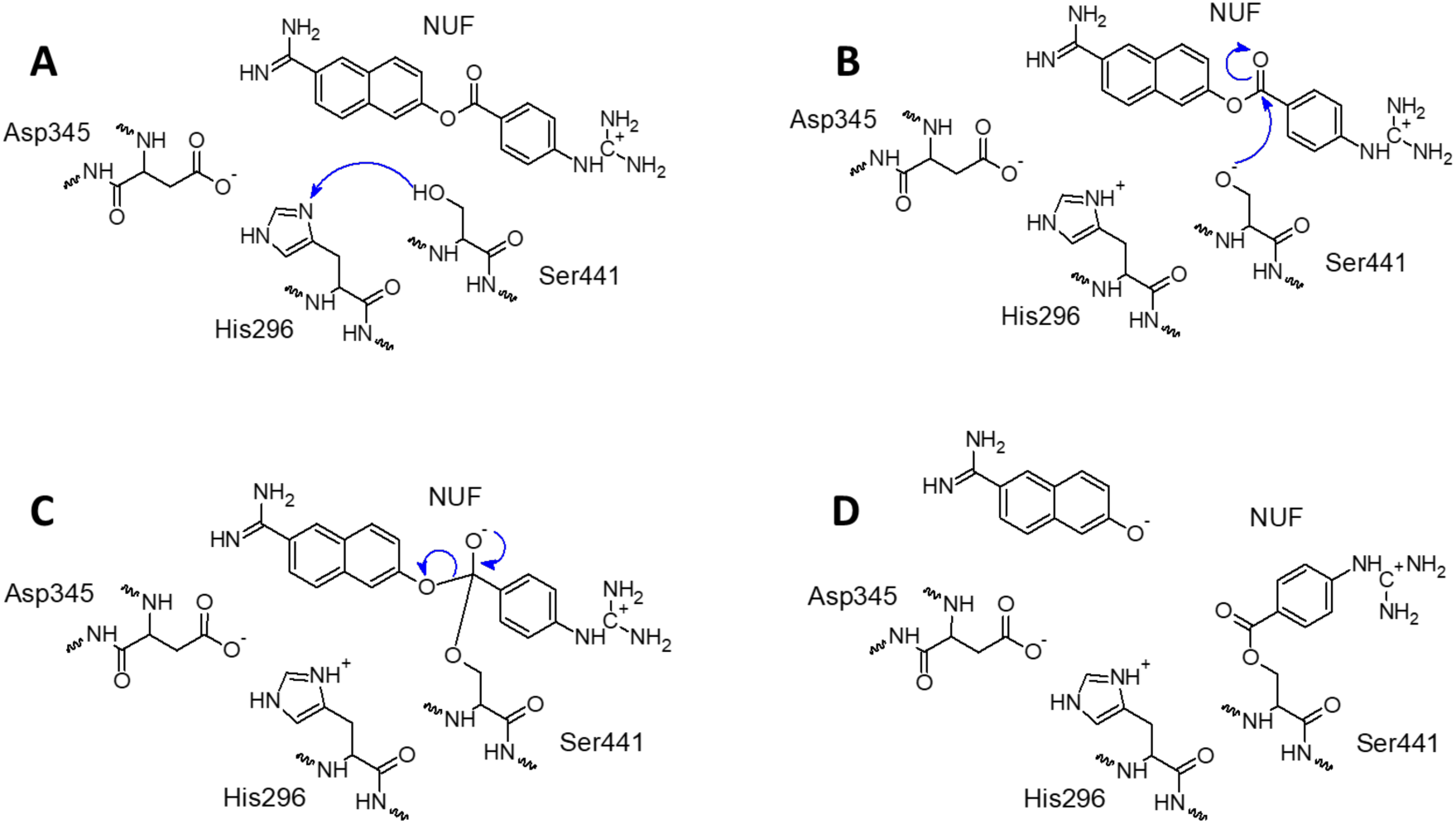
Reaction mechanisms for the transesterification of NUF. A) reactive, B) transition state, C) intermediate, D) product

By using a fully multiscale approach, combining classical MD simulation with enhanced sampling QM/MM approaches, we have unraveled the structural features of the interaction of TMPRSS2 with a lipid bilayer. In particular, we have identified the stability of the membrane protein complex, and we have spotted out the formation of salt-bridges in the vicinity of the active site upon the cleavage at the Arg255-Ile256 position, leading to the enzyme maturation. We have also examined the interaction between TMPRSS2 and its proposed inhibitor NUF. While the non-covalent complex appears labile, the irreversible poisoning of the catalytic Ser441 involves only a moderate energy barrier. This fact is also due to the presence of His296 which is capable of activating the serine by deprotonating its terminal -OH group, bypassing a negligible barrier of only 3 kcal/mol.

The covalent inhibition of TMPRSS2 is a most attractive therapeutic strategy, due to the involvement of the enzyme in both cancer progression and viral reproduction, including SARS-CoV-2. If the chemical reactivity leading to the serine-poisoning appears already highly optimized, increasing the affinity of the non-covalent complex by favoring specific interactions in addition to the dominant hydrophobic pattern could be mostly beneficial to boost inhibition. Furthermore, identifying a better leaving group, as for instance a more stabilized alcoholate, could increase the reaction driving force, further contributing to increasing the inhibition power.

Globally, we have shown that our state-of-the-art multiscale approach and high-level molecular modeling and simulation are able to provide a fully resolved picture of the functioning of complex enzymes and their inhibition by external agents, hence, constituting a most valuable asset for rational drug design.

## ASSOCIATED CONTENT

### Supporting Information

Extended computational methodology. Main docking poses for the NUF/TMPRSS2 complex; identification of the sulfur bridges stabilizing TMPRSS2 conformation; simulation of non-sulfur bonded TMPRSSS2; analysis of the convergence of the String method and decomposition of the reaction free energy on individual coordinates; snapshots around critical points of the reaction PMF; NUF force field parameters.

(file type, i.e., PDF)

## ACKNOWLEDGMENT

All the simulations have been performed on the LPCT, Explor, and GENCI computing resources which are gratefully acknowledged. Financial support from the French Ministry for Higher Education and Research (MESR) and CNRS is also acknowledged. A.M. and F.B. thank ANR and CGI for their financial support of this work through Labex SEAM ANR 11 LABX 086, ANR 11 IDEX 05 02. The support of the IdEx “Université Paris 2019” ANR-18-IDEX-0001. A.S. and G.B. thanks for financial support the European Union - NextGenerationEU through the Italian Ministry of University and Research under PNRR - M4C2-I1.3 Project PE_00000019 “HEAL ITALIA” CUP (B73C22001250006).

