## Supplementary Information for "Mechanism of the Covalent Inhibition of Human Transmembrane Protease Serine 2 as an Original Antiviral Strategy"

**Corresponding Author**

EXTENDED COMPUTATIONAL METHODOLOGY

*Equilibrium MD simulation*

Two different initial systems have been constructed starting from the experimental pdb:7meq file of TMPRSS2 (see main text).[1] In one instance the disulfide bridges have been enforced by deprotonating the corresponding cysteine residues and adding an extra bond between the two nearby SG atoms via the amber *tleap* utility. In a second system the cysteine residues have been kept protonated and no extra bond was added. As expected, the second system shows a higher flexibility confirming the importance of the sulfur bridge network in the conformational stability of TMPRSS2. The equilibrium MD simulation lasted for 500 ns.

The protein and the POPC lipids have been modeled using the amberff14 force field,[2] while water is represented with the TIP3P force field.[3] The Newton equations of motions have been propagated with a time step of 4 fs, thanks to the combined use of Rattle and Shake[4] and the Hydrogen Mass Repartition (HMR) approach.[5] The equilibration and thermalization was achieved by progressively relaxing positional constraints on the lipid and protein heavy atoms, and the MD simulation was performed in the isobaric and isothermal ensemble (NPT) at 300 K and 1 atm. The Langevin thermostat and barostat have been consistently used.

The same procedure has been used for the equilibrium MD simulations of the system after maturation, i.e. the proteolytic cleavage of the Arg255/Ile256 bond, and after the docking with NUF. As for the ligand the force field has been parameterized via the Generalized Amber Force Field (GAFF) strategy.[6] Notably point charges have been obtained fitting the restricted electrostatic potential (RESP) calculated form the equilibrium geometry of the ligand at HF/6-31G level of theory, as standard.

*Steered MD Simulation*

After proteolytic cleavage, Ile256 terminal group remains locked in a metastable position due to the electrostatic and steric hindrances of other nearby amino acids. Thus, the formation of the salt-bridge with Asp440 cannot be spontaneously observed. Consequently, we have performed steered MD simulation (SMD)[7] forcing the distance between the centers of mass of the two amino acids to decrease from the initial value of 13.8 to 3.3 Å. To this aim a harmonic potential of 5.0 kcal/mol was applied and its center displaced progressively during 1,000,000 steps.

At the end of the SMD run, the harmonic potential was lifted and an equilibrium MD simulation was propagated for 700 ns.

To avoid a computational overload and possible string instability, a reactive conformation between TMPRSS2 catalytic triad and NUF was achieved implementing again classical SDM. In this case the distance between the centers of mass of NUF and the catalytic triad (His296, Asp345, and Ser441) was reduced from 5.0 to 2.0 Å in 100,000 steps.

*Docking*

To correctly place the NUF ligand inside the binding site of TMPRSS2, molecular docking was performed. The binding site, which was centered on the catalytic Serine, was represented as a box of 22x18x21 $Å^3$ in the x, y, and z directions. Three docking software, namely Autodock[8], Vina[9], and Smina[10], were utilized for this task we the same binding box. The best docking pose, which is presented in figure S0 for all three software, was found to be highly similar, thereby emphasizing the convergence of the results. Subsequently, the docking position of NUF was selected for further molecular dynamics (MD) simulations.

*Umbrella Sampling*
The umbrella sampling (US) technique[11] based on classical MD simulation was employed to determine the free energy for the non-covalent association of NUF with TMPRSS2. The chosen collective variable used to describe the process was the distance between the heavy atoms of NUF and the side chains of TMPRSS2 catalytic triad, i.e. His296, Asp345 and Ser441. A weak harmonic restraint of 5 kcal/mol.Å² was imposed on the collective variable, which started from a distance of 11.5 Å and was gradually increased to 26.5 Å in steps of 0.25 Å. As a result, 60 independent biased MD trajectories, each with a duration of 50 ns, were conducted. It should be noted that the force acting on the NUF ligand was applied to the x and y directions, only. Since, the membrane is roughly oriented in the (xOy) plane, this choice, which leaves unbiased the distance between NUF and the lipid bilayer, enables to differentiate the situations in which the ligand is either fully solvated, or interacting with the membrane polar heads. The final 40 ns of the 60 overlapping trajectories were used for determining the potential of mean force (PMF) with the weighed histogram analysis method[12].

*Reactivity and String Method*
To assess the covalent inhibition of TMPRSS2 by NUF we resorted to semiempirical QM/MM methods at the DFTB3 level of theory (see main text). The QM partition comprised the full NUF, and the side chains of the amino acids constituting the catalytic triad (His296, Asp345, and Ser441). The dangling C$\alpha$-C$\beta$ bond was treated using the link atom approach. QM/MM was performed using the Amber code, a time step of 1 fs was used for propagating the equations of motion. An initial equilibrated structure of the reactant was obtained through QM/MM equilibrium starting from a reactive conformation obtained by classical MD. The product state was obtained performing SMD at semiempirical QM/MM level and notably enforcing the breaking of the CO bond in NUF, the transfer of the proton of Ser441 OH to the $\varepsilon$ N atom of His296, and the formation of the bond between Ser441 and the carbonyl of NUF. As detailed in the main text initial strings were generated considering a linear combination of the distances between the carbon and oxygen atoms in NUF, the NUF carbon atom and the lateral oxygen of Ser441, and between the hydrogen atom of Ser441 and the N$\varepsilon$ atom of His296.
The string has been optimized with the adaptive procedure[13] for 20,000 steps corresponding to 20 ps. After having defined the optimized minimum energy path, replica exchange umbrella sampling on the string space has been performed for a total of 320,000 steps (320 ps). The maximum statistical error, defined as the 95% probability, has been estimated to 0.6 kcal/mol, while the incertitude amounts at 0.3 kcal/mol at the transition state.
As reported in Figure S3, the full PMF has also been decomposed on the contribution of the individual collective variables.

*References*

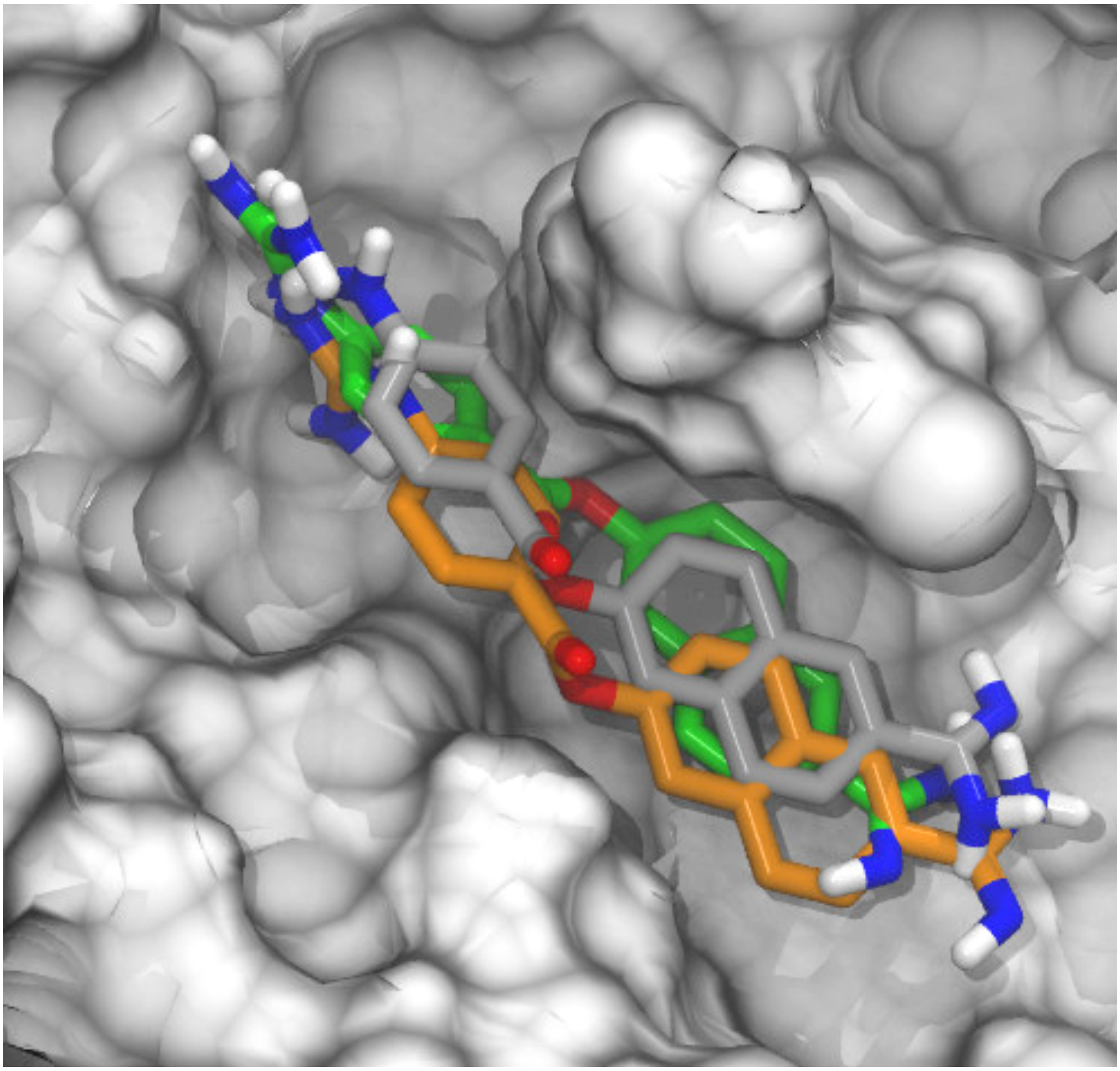

Figure S1: The three most favorable molecular docking poses obtained for Autodock4 (carbon green), SMINA (carbon grey) and VINA (carbon orange).

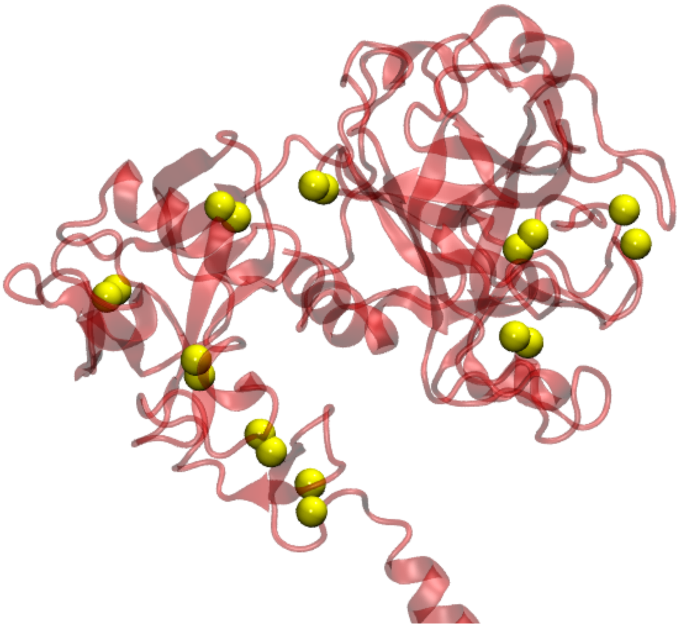

Figure S2. Representation of the SG atom (yellow van der Waals spheres) of the cysteine residues involved in the disulfide-bridge network in TMPRSS2.

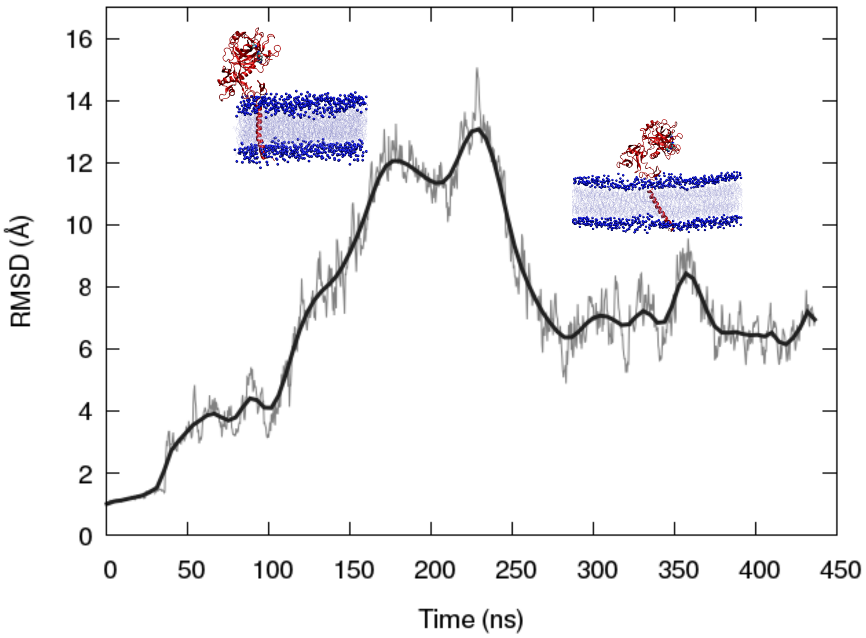


Figure S3. Time evolution of the RMSD for the membrane embedded TMPRSS2 without enforcing sulfur bonds, showing the higher instability of the protein. Two representative snapshots are given in the inlay.

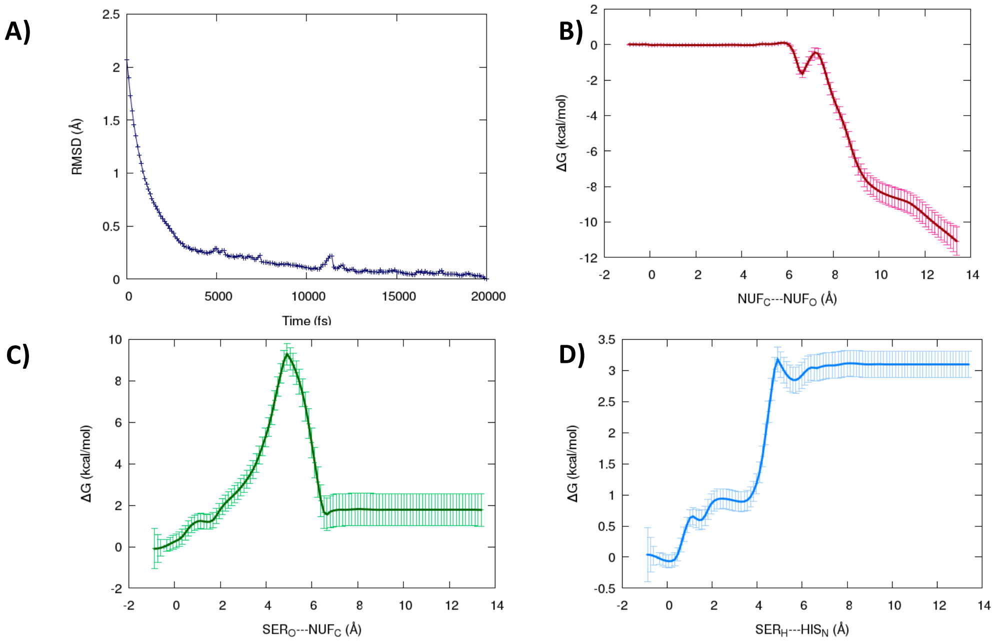


Figure S4. Convergence of the string optimization process by monitoring the RMSD evolution. Decomposition of the PMF into the contribution due to the individual collective variables (B-D) the same color code as in Figure 5 of the main text is used to differentiate the coordinates.

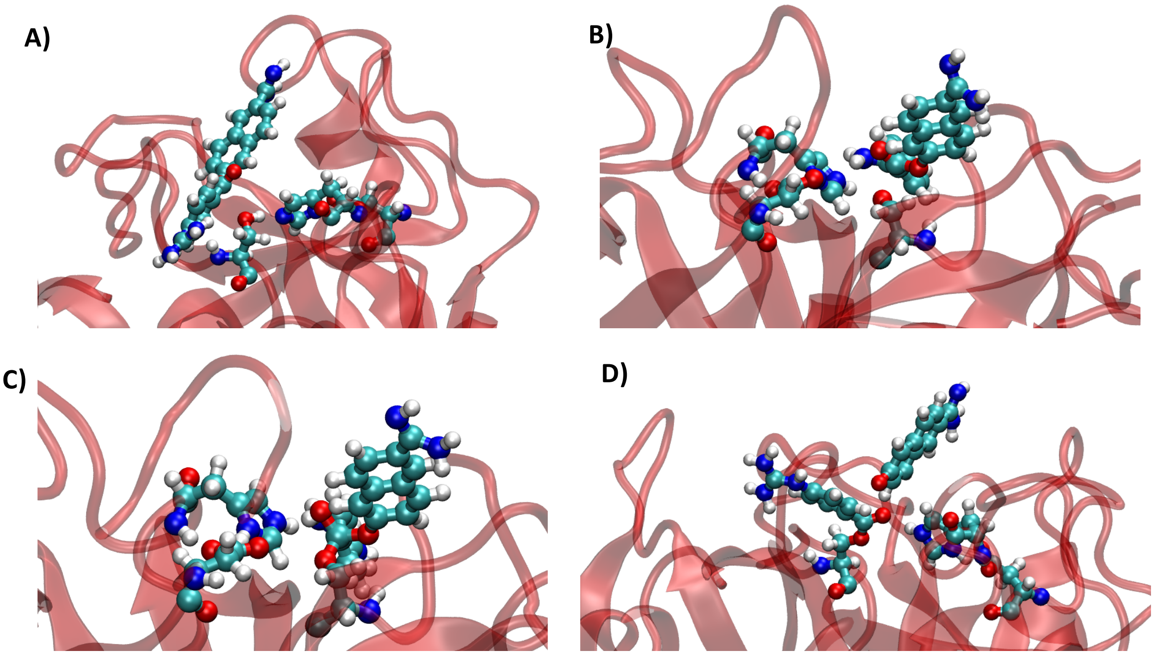


Figure S6. Representative snapshots illustrating the structure corresponding to the critical points obtained along the PMF A) reactive, B) transition state, C) intermediate, D) product.

**Equilibrium Geometry, atom types and RESP charges of NUF**

| | | | | | | | |
|---|---|---|---|---|---|---|---|
| 1 | C1 | 7.7240 | -0.1070 | -0.1670 | CA | 1 NUF | 0.999592 |
| 2 | N1 | 7.6310 | 1.2190 | -0.0820 | N2 | 1 NUF | -0.968536 |
| 3 | N2 | 8.9170 | -0.7040 | -0.1660 | N2 | 1 NUF | -0.968536 |
| 4 | N3 | 6.6190 | -0.8800 | -0.2620 | N2 | 1 NUF | -0.689046 |
| 5 | H1 | 6.7650 | -1.8140 | -0.6270 | H | 1 NUF | 0.407555 |
| 6 | C2 | 5.2640 | -0.4910 | -0.0730 | CA | 1 NUF | 0.354224 |
| 7 | C3 | 4.8800 | 0.3630 | 0.9700 | CA | 1 NUF | -0.273834 |
| 8 | C4 | 3.5370 | 0.6830 | 1.1300 | CA | 1 NUF | -0.086351 |
| 9 | C5 | 2.5660 | 0.1430 | 0.2760 | CA | 1 NUF | -0.117923 |
| 10 | C6 | 2.9580 | -0.7280 | -0.7510 | CA | 1 NUF | -0.086351 |
| 11 | C7 | 4.3010 | -1.0380 | -0.9300 | CA | 1 NUF | -0.273834 |
| 12 | C8 | 1.1470 | 0.5160 | 0.5090 | C | 1 NUF | 0.799123 |
| 13 | O1 | 0.7630 | 1.2840 | 1.3670 | O | 1 NUF | -0.544766 |
| 14 | O2 | 0.3080 | -0.1210 | -0.3600 | OS | 1 NUF | -0.423235 |
| 15 | C9 | -1.0650 | 0.1440 | -0.2920 | CA | 1 NUF | 0.318604 |
| 16 | C10 | -1.5530 | 1.4270 | -0.6360 | CA | 1 NUF | -0.253056 |
| 17 | C11 | -2.9100 | 1.6460 | -0.6480 | CA | 1 NUF | -0.244881 |
| 18 | C12 | -3.8250 | 0.6030 | -0.3280 | CA | 1 NUF | 0.203653 |
| 19 | C13 | -3.3100 | -0.6900 | 0.0160 | CA | 1 NUF | 0.134561 |
| 20 | C14 | -1.9050 | -0.8940 | 0.0250 | CA | 1 NUF | -0.319199 |
| 21 | C15 | -5.2280 | 0.8040 | -0.3390 | CA | 1 NUF | -0.273571 |
| 22 | C16 | -6.1050 | -0.2170 | -0.0220 | CA | 1 NUF | -0.112694 |
| 23 | C17 | -5.5840 | -1.4950 | 0.3280 | CA | 1 NUF | -0.068986 |
| 24 | C18 | -4.2280 | -1.7250 | 0.3390 | CA | 1 NUF | -0.263784 |
| 25 | C19 | -7.5820 | 0.0280 | -0.0530 | CM | 1 NUF | 0.748185 |
| 26 | N4 | -8.1930 | 0.8080 | -0.8790 | NC | 1 NUF | -0.841760 |
| 27 | N5 | -8.3160 | -0.7020 | 0.8630 | NT | 1 NUF | -0.938885 |
| 28 | H2 | 8.4480 | 1.7900 | 0.0880 | H | 1 NUF | 0.472478 |
| 29 | H3 | 6.7670 | 1.6940 | -0.3040 | H | 1 NUF | 0.472478 |
| 30 | H4 | 9.7700 | -0.1700 | -0.2520 | H | 1 NUF | 0.472478 |
| 31 | H5 | 9.0050 | -1.7020 | -0.0330 | H | 1 NUF | 0.472478 |
| 32 | H6 | 5.6160 | 0.7540 | 1.6640 | HA | 1 NUF | 0.189565 |
| 33 | H7 | 3.2270 | 1.3430 | 1.9330 | HA | 1 NUF | 0.176176 |
| 34 | H8 | 2.2150 | -1.1550 | -1.4140 | HA | 1 NUF | 0.176176 |
| 35 | H9 | 4.6050 | -1.7000 | -1.7350 | HA | 1 NUF | 0.189565 |
| 36 | H10 | -0.8550 | 2.2160 | -0.8900 | HA | 1 NUF | 0.183202 |
| 37 | H11 | -3.3000 | 2.6250 | -0.9120 | HA | 1 NUF | 0.193151 |
| 38 | H12 | -1.4940 | -1.8670 | 0.2780 | HA | 1 NUF | 0.189206 |
| 39 | H13 | -5.6100 | 1.7920 | -0.5840 | HA | 1 NUF | 0.157543 |
| 40 | H14 | -6.2720 | -2.3000 | 0.5660 | HA | 1 NUF | 0.131406 |
| 41 | H15 | -3.8430 | -2.7080 | 0.5950 | HA | 1 NUF | 0.171947 |
| 42 | H16 | -7.5300 | 1.1720 | -1.5650 | H | 1 NUF | 0.351460 |
| 43 | H17 | -7.8820 | -0.8820 | 1.7600 | H | 1 NUF | 0.392211 |
| 44 | H18 | -9.2850 | -0.4140 | 0.9350 | H | 1 NUF | 0.392211 |
